# Analytical expressions and physical principles for single-cell mRNA distributions of the *lac* operon of *Escherichia coli*

**DOI:** 10.1101/520478

**Authors:** Krishna Choudhary, Atul Narang

## Abstract

Mechanistic models of stochastic gene expression are of considerable interest, but their complexity often precludes tractable analytical expressions for mRNA and protein distributions. The *lac* operon of *E. coli* is a model system with regulatory elements such as multiple operators and DNA looping that are shared by many operons. Although this system is complex, intuition suggests that fast DNA looping may simplify it by causing the repressor-bound states of the operon to equilibrate rapidly, thus ensuring that the subsequent dynamics are governed by slow transitions between the repressor-free and the equilibrated repressor-bound states. Here, we show that this intuition is correct by applying singular perturbation theory to a mechanistic model of *lac* transcription with the scaled time constant of DNA looping as the perturbation parameter. We find that at steady state, the repressor-bound states satisfy detailed balance and are dominated by the looped states; moreover, the interaction between the repressor-free and the equilibrated repressor-bound states is described by an extension of the Peccoud-Ycart two-state model in which both (repressor-free and repressor-bound) states support transcription. The solution of this extended two-state model reveals that the steady state mRNA distribution is a mixture of the Poisson and negative hypergeometric distributions which reflects mRNAs obtained by transcription from the repressor-bound and repressor-free states, respectively. Finally, we show that the physics revealed by perturbation theory makes it easy to derive the extended two-state model equations for complex regulatory architectures.

## 1 INTRODUCTION

Stochastic models of single-cell gene expression have become progressively more complex. The earliest models had no regulation since they assumed that the promoter and ribosome binding site were always available for initiation, and stochasticity arose from the random initiation of transcription and translation (1–3). These *one-state models* were followed by *two-state models* in which the promoter switched randomly between two states of which only one was available for initiation of transcription (4–6). Rigorous comparison of stochastic models with experimental data requires analytical expressions for the distributions of mRNA and protein, rather than their mean and variance (6). To facilitate such comparison, analytical expressions for the steady state mRNA and protein distributions have been derived for the one-state and two-state models (1, 4–10).

Early experiments with unregulated promoters, which used GFP intensity as a reporter for protein number, were consistent with the one-state model (11, 12). However, subsequent experiments using single-molecule techniques have led to considerable debate regarding the origin of the stochasticity (13, 14). Specifically, Choi *et al.* measured the steady state distributions of LacY obtained when the *lac* operon of *Escherichia coli* is uninduced or partially induced. They observed that the size, but not the frequency, of large protein bursts increased dramatically in partially induced cells, and explained this result in terms of specific molecular mechanisms known to operate in the *lac* operon of *E. coli.* In contrast, So *et al.* found that essentially the same noise was observed in several operons of *E. coli* regulated by very different molecular mechanisms. Thus, the central question of this debate is whether the noise arises from gene-specific or genome-wide mechanisms (15–18).

Attempts to resolve the foregoing question have led to several mechanistic models of stochastic gene expression taking due account of the molecular mechanisms that regulate gene expression. Some have focused specifically on models of *lac*expression in *E. coli*(19–22), while others have studied generic models spanning a wide range of regulatory mechanisms (23). However, in all these models, the operon can exist in more than two states. To investigate steady state mRNA distributions from them, one must work with a system of non-autonomous first-order ordinary differential equations of dimension equal to the number of states of the operon. Study of such systems has been done using analytical expressions for the first two moments of the mRNA distribution or, expressions for complete distributions using the so-called binomial moment method (22–26). However, these methods result in quite intricate and intractable expressions. Alternatively, one determines the mRNA distribution computationally rather than analytically (19–21).

Even in mechanistic models of gene expression, there are disparate time scales that can be exploited. In the particular case of the *lac* operon, DNA looping is much faster than repressor-DNA binding or dissociation (21). In earlier work (27), which was aimed at gaining deeper insights into the data of Choi *et al*., we exploited the existence of fast DNA looping to derive a simple analytical expression for the steady state protein distribution in the uninduced and partially induced *lac* operon of *E. coli.*This expression was obtained by arguing heuristically that at steady state, the looped states are dominant, the repressor-bound species satisfy detailed balance, and the model reduces to the *leaky two-state model,* an extension of the two-state model in which the repressor-bound states also permit transcription at a low rate. We confirmed the validity of our heuristic approach by showing that our analytical expressions agreed well with stochastic simulations of the model, but we did not provide rigorous justification for our results.

It turns out that there is a well-developed singular perturbation theory for systematically simplifying stochastic models with disparate time scales (28, 29). Recently, this theory and related quasi-steady state approximations have been utilized successfully to study single-cell gene expression noise (30, 31). Here, we use singular perturbation theory to analyze a stochastic model for transcription of the uninduced or partially induced *lac* operon of *E. coli*. We show that the results obtained, with the scaled time constant of DNA looping as the perturbation parameter, are identical to those obtained from our heuristic approach (27). The zeroth-order solution shows that the repressor-bound states are predominantly in the looped state. The solution to first-order shows that at steady state, the repressor-bound species satisfy detailed balance, and the interactions of the repressor-bound and repressor-free species are described by leaky two-state model. Finally, the theory yields simple and physically meaningful expressions for the mean, variance, and generating function of the steady mRNA distribution. In addition, we show that alternative parameter regimes, e.g., operator-repressor dissociation rate being comparable to DNA looping rate, may also result in the leaky two-state model.

## 2 MODEL

### 2.1 Description

The regulation of the *lac* operon involves a single transcription factor and multiple operators (32, 33). The transcription factor is a repressor protein denoted R which can bind specifically to three sites on the chromosome, namely the main operator *O*_1_ and two auxiliary operators, *O*_2_ and *O*_3_. Although we shall consider this problem later on, it is convenient to begin by considering the special case of only one auxiliary operator *O*_2_ in which case the *lac* operon can exist in only 4 states (Fig.1). If both operators are free of repressor, the operon is in the *repressor-free state* denoted O. If a repressor binds either *O*_1_ or *O*_2_, the operon contains a single repressor; we denote these states by *O*_1_ · *R* and *O*_2_ · *R*. Finally, since the repressor is a tetramer, it can bind both operators simultaneously leading to the formation of a looped state denoted *O*_1_ · *R* · *O*_2_. In principle, two repressors can also bind to each of the two operators sites yielding an operon with two repressors, but experiments show that in wild-type cells, the likelihood of such multi-repressor operons is negligibly small. Thus, according to our model, the *lac* operon can only be in one of the four states, namely the repressor-free state O and the three *repressor-bound states O*_1_ · *R*, *O*_2_ · *R*, and *O*_1_ · *R* · *O*_2_.

**Figure 1:**
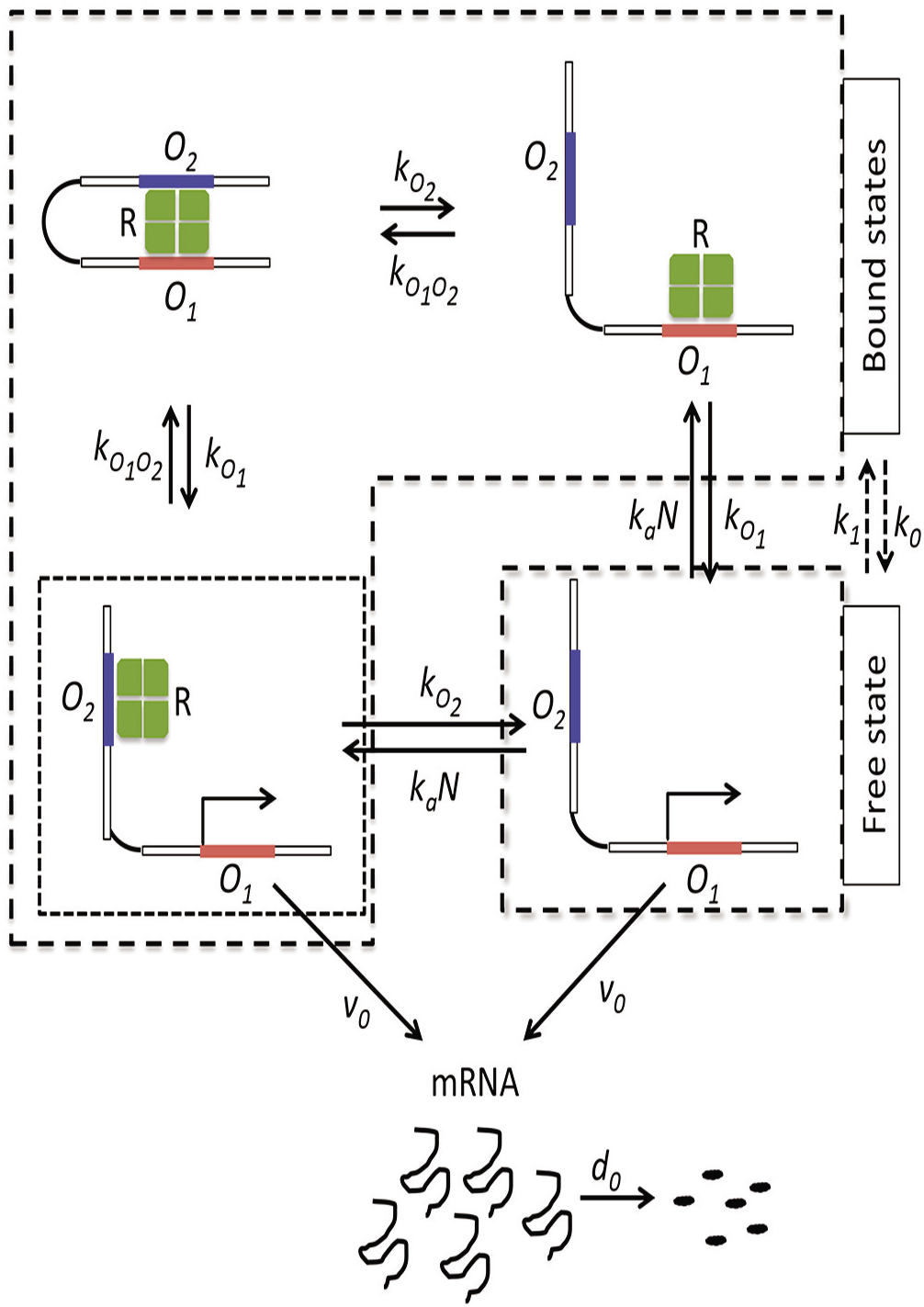
The model scheme: The *lac* operon is assumed to have four states: A repressor-free state, denoted *O*, in which both the main operator *O*_1_ and the auxiliary operator *O*_2_ are free of repressor; two non-looped repressor-bound states, *O*_1_ · *R* and *O*_2_ · *R*, in which the repressor is bound to *O*_1_ and *O*_2_, respectively; and a looped repressor-bound state *O*_1_ · *R* · *O*_2_ in which the repressor is simultaneously bound to *O*_1_ and *O*_2_.

The foregoing four states inter-convert with the propensities shown in Fig.1. Since binding of cytosolic repressor to an operator is diffusion-limited, it occurs with the same propensity *k_a_N* regardless of the operator, where *k_a_* is the propensity for association of a single cytosolic repressor to an operator, and N is the number of repressors per cell (assumed to be the same for all cells of the population). The propensity for dissociation of a repressor from an operator does depend on the identity of the operator; we denote the propensities for dissociation from *O*_1_ and *O*_2_ by *k*_*O*_1__ and *k*_*O*_2__, respectively. If the repressor is bound to *O*_1_ or *O*_2_, it can form the looped state *O*_1_ · *R* · *O*_2_ by binding to the other operator with propensity *k*_*O*_1_*O*_2__. Of the four states, only *O* and *O*_2_ · *R* allow transcription, and transcription from both states occurs with the same propensity vo. Finally, the mRNA degrades with propensity *d*_0_.

### 2.2 Master equations

We model the kinetic scheme shown in Fig.1 with chemical master equations. The instantaneous state of the system is described by two variables, namely the mRNA copy number m, and the state of the operon s where s is chosen as *f*, 1, 2, and 12 when the operon is in the states *O, O*_1_ · *R, O*_2_ · *R* and *O*_1_ · *R* · *O*_2_, respectively. We denote the instantaneous and steady state probabilities of *m* mRNAs when the operon is in the state *s* by *p_m,s_* and 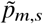, respectively. The master equations for the model are

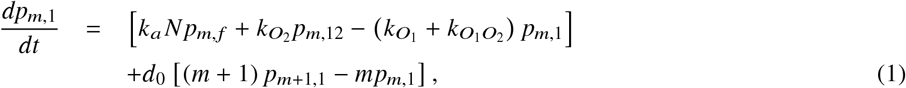

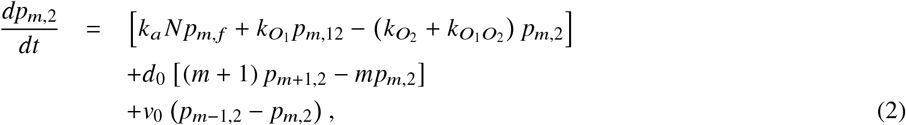

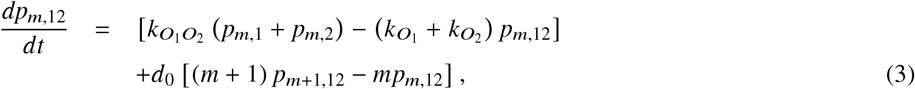

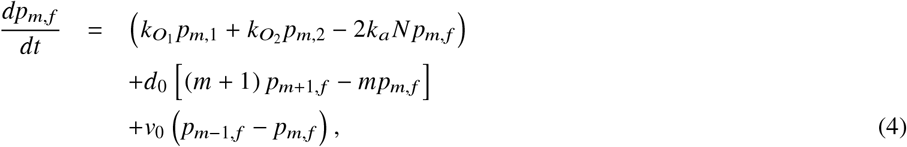

and the normalization condition is

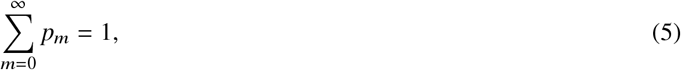

where

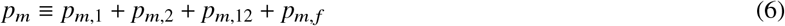

is the marginal mRNA distribution. Our goal is to determine the steady state marginal mRNA distribution 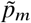.

**Table 1:** Parameter values in the absence of inducer (27).

| Parameter | Value (in $s^{-1}$) | Parameter | Value (in $s^{-1}$) |
| --- | --- | --- | --- |
| $k_{O_1}$ | 0.0016 | $k_a N$ | 0.07 |
| $k_{O_2}$ | 0.019 | $v_0$ | 0.12 |
| $k_{O_1 O_2}$ | 4 | $d_0$ | 0.011 |

## 3 RESULTS

### 3.1 Setting up equations for application of singular perturbation theory

Singular perturbation theory exploits the existence of disparate time scales. In the *lac* operon, the looping propensity *k*_*O*1*O*2_ is much greater than the propensity for any other reaction (Table 1). It is therefore conceivable that the repressor-bound states undergo looping so rapidly that they equilibrate on the fast time scale 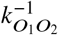, after which there are relatively slow transitions between the repressor-free and repressor-bound states. To capture this physical argument, it is convenient to replace one of the original fast variables *p*_*m*,1_, *p*_*m*,2_, *p*_*m*,12_ with the slow variable

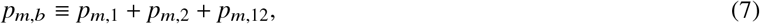

which represents the probability of *m* mRNAs when the operon is repressor-bound. If we replace *p_m_*,_12_ with *p_m,b_*, Eq. 4 remains unchanged, but the evolution of the repressor-bound states is given by the new equations

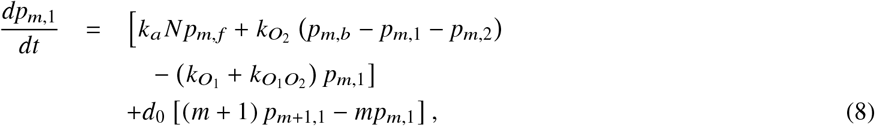

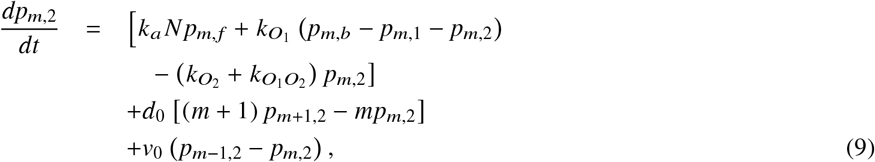

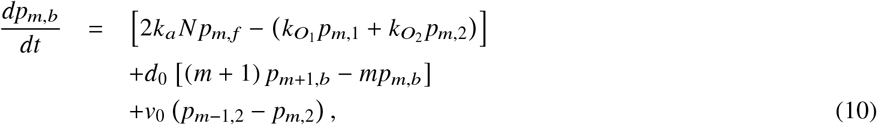

and the normalization condition (Eq. 5) becomes

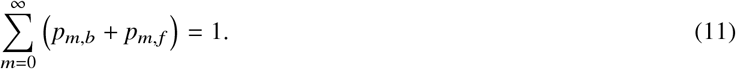

Henceforth, we shall work with Eq. 4, 8–11.

We apply singular perturbation theory to these equations by following the method developed by Rawlings and coworkers (28, 29). Multiplying Eq. 4 and Eq. 8–10 with the perturbation parameter 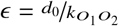, and defining 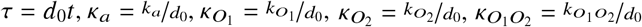 and 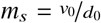 gives the slow equations

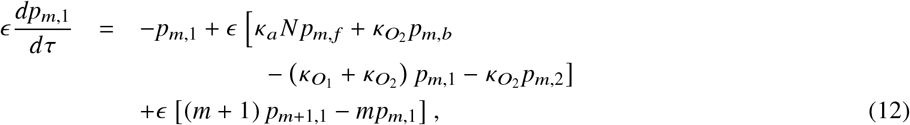

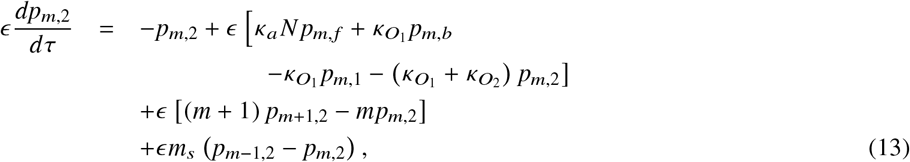

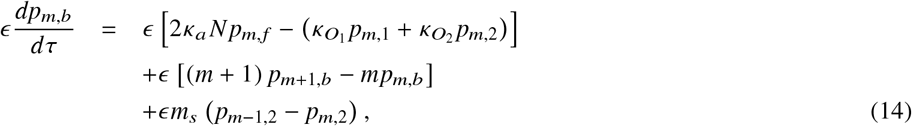

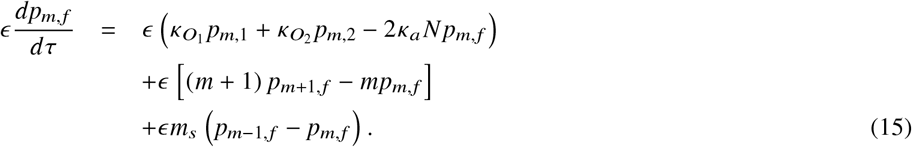

Next we substitute the power series expansions

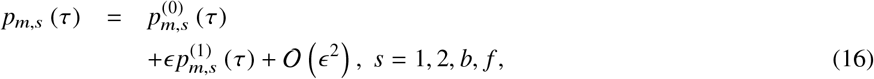

in Eq. 11–15, and collect terms with coefficients *ϵ*^0^ and *ϵ*^1^ to determine the zeroth- and first-order relations.

### 3.2 Zeroth-order equations imply that the operon is completely repressed at steady state

Collecting terms with coefficient *ϵ*^0^ yields the zeroth-order equations

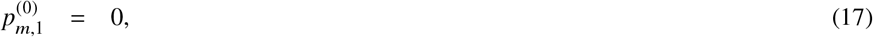

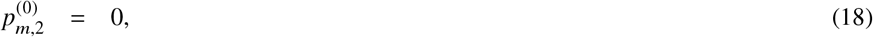

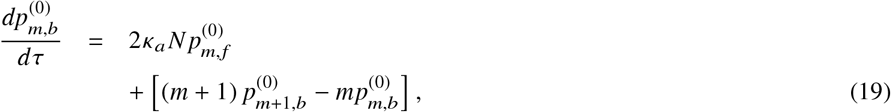

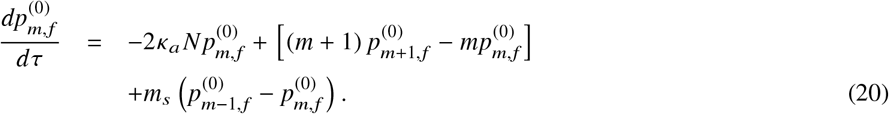

and the zeroth-order normalization condition

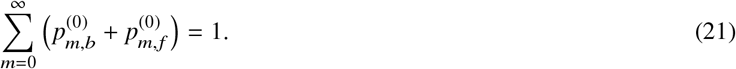

It follows from Eq. 17–18 that to zeroth order, the probabilities of the *O*_1_ · *R* and *O*_2_ · *R* states are always zero, and the repressor-bound species are always exclusively in the looped state *O*_1_ · *R* · *O*_2_, i.e., 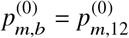.

Eq. 19–21 immediately yield the steady state probabilities 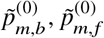. To see this, observe that Eq. 21 implies that 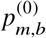 and 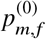 behave like probabilities. The evolution of these probabilities is governed by Eq. 19–20, which are formally similar to the equations of the two-state model, the only difference being that there is a *unidirectional* probability flux 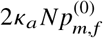 from the repressor-free to the repressor-bound state. It follows that ultimately, 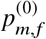 approaches zero, i.e., 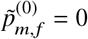 for all *m* ≥ 0, and the operon is in the repressor-bound state, i.e, 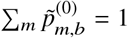. Now since 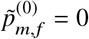, there is no transcription from the repressor-free state *O* at steady state, and since 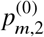 is always zero, there is no transcription from the state *O*_2_ · *R* either. It follows that there is no mRNA at steady state, i.e, 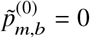 for all *m* ≥ 1, and the zeroth-order steady state probabilities are

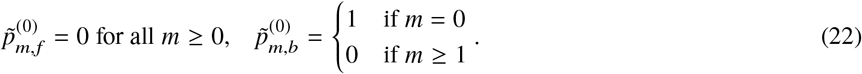

This is shown rigorously in Appendix A by solving Eq. 19–21 at steady state by the method of generating functions.

### 3.3 Detailed balance of bound species and reduction to leaky two-state model at steady state

Collecting first-order terms yields the first-order equations

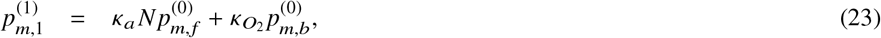

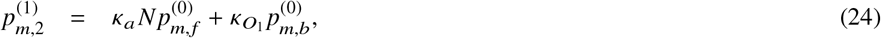

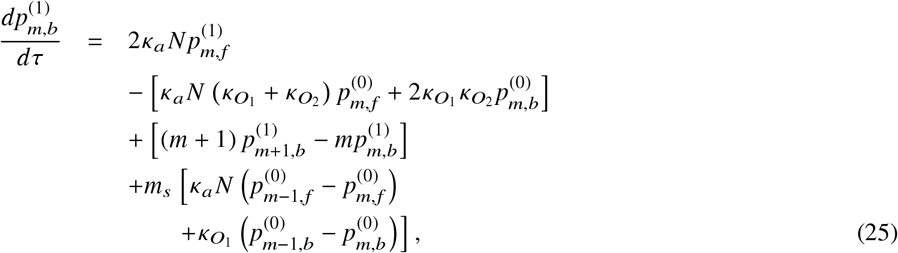

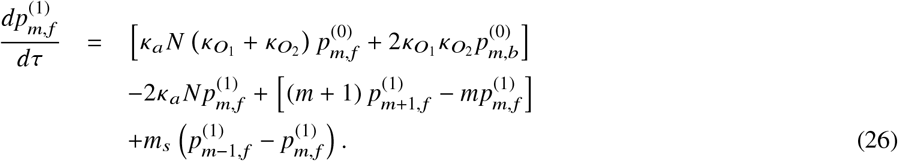

and the first-order normalization condition

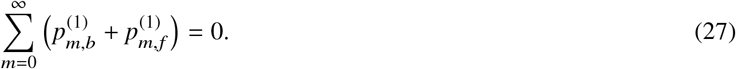

It is clear from Eq. 27 that 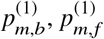 are not probabilities, and hence Eq. 25–26 do not lend themselves to simple physical interpretation. However, we show below that at steady state, the zeroth- and first-order solutions, taken together, imply that the repressor-bound species satisfy detailed balance, and the interaction between the repressor-free and repressor-bound species is described by the leaky two-state model.

At steady state, Eq. 22 holds, and Eq. 23–26 reduce to the simpler form

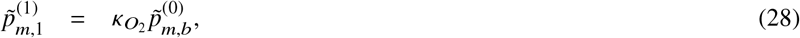

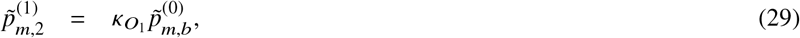

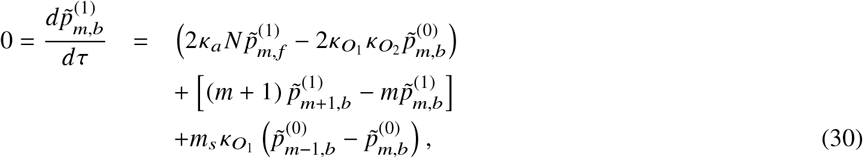

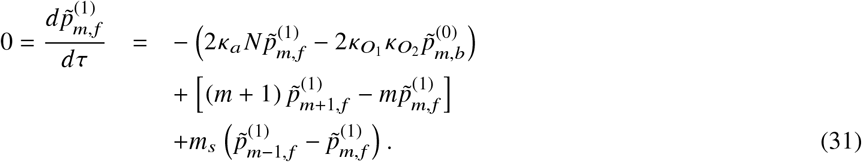

These equations can be solved for 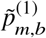 and 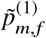 by the method of generating functions (Appendix A), but we shall focus hereafter on the physics revealed by these equations.

Eq. 28–29 imply that the repressor-bound states satisfy the principle of detailed balance to order *ϵ*. Indeed, multiplying Eq. 28–29 by e and appealing to the zeroth-order relations 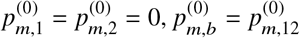 yields the equations

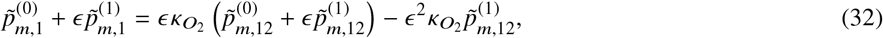

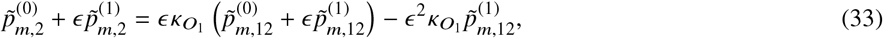

which imply that 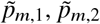, and 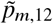 satisfy the detailed balanced equations 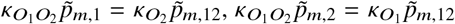 to order *ϵ*. Note that application of the same argument to Eq. 28–29, but without replacing 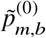 with 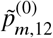, shows that to order *ϵ*, we also have the relations 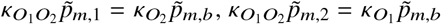, and the marginal probabilities of the *O*_1_ · *R* and *O*_2_ · *R* states are 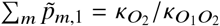 and 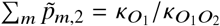, respectively. We shall use these relations later.

Eq. 30–31 imply that to order *ϵ*, the steady state probabilities of the repressor-bound and repressor-free states 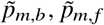 are described by the leaky two-state model. Indeed, multiplying Eq. 30–31 by e and appealing to the zeroth-order solutions (Eq. 22) yields the relations

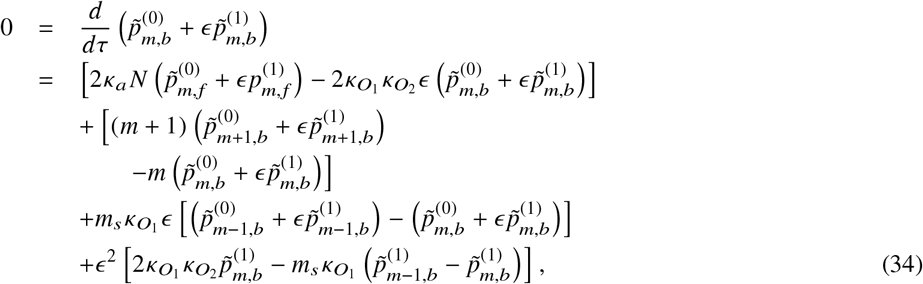

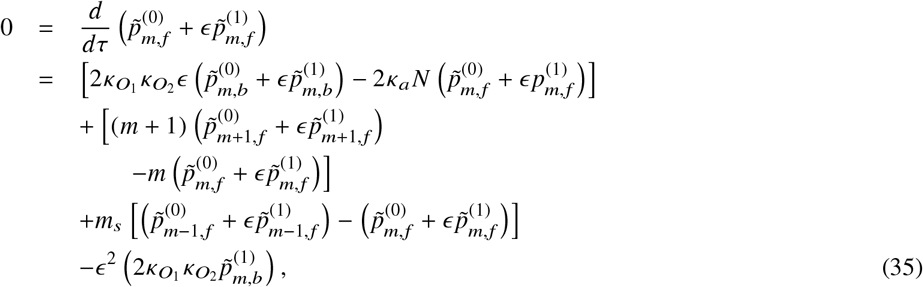

which imply that to order *ϵ*, the steady state probabilities 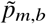 and 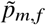 satisfy the equations

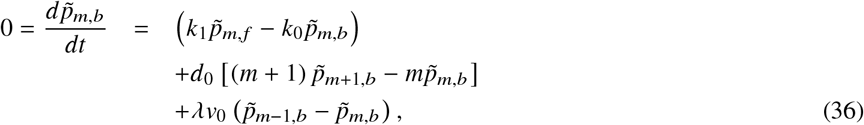

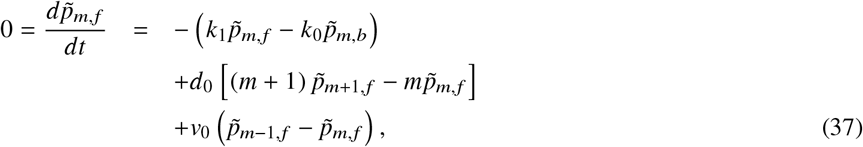

where

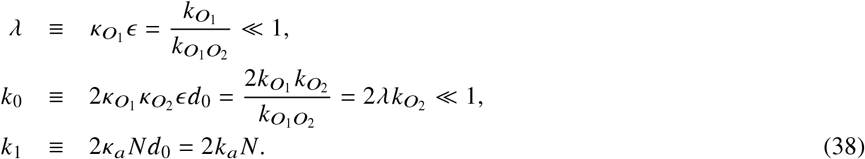

Eq. 36–38 describe the steady states of the *leaky two-state model* (Fig. 2) in which transitions occur between repressor-bound and repressor-free states with propensities *k_0_* and *k*_1_, but the repressor-bound state also allows *leaky* transcription with a low propensity *λν*_0_.

**Figure 2:**
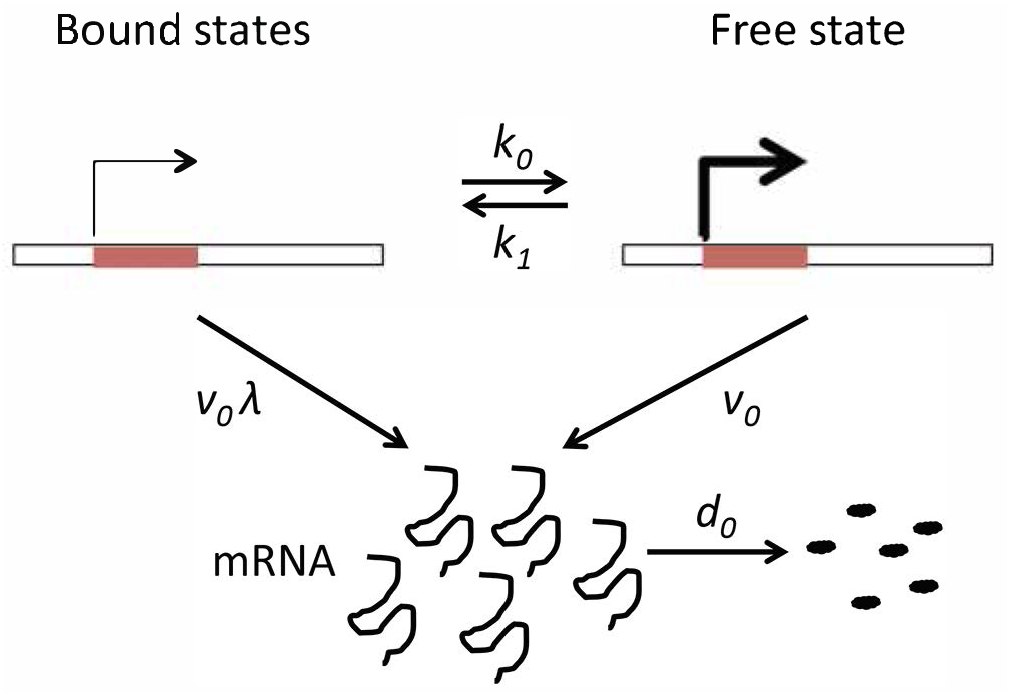
The leaky two-state model. Like the two-state model, the operon switches randomly between the repressor-free and the repressor-bound states with propensities *k*_0_, *k*_1_, and the repressor-free state permits transcription with propensity *ν*_0_. However, unlike the two-state model, the leaky two-state model also permits transcription from the repressor-bound state with a small propensity *λν*_0_.

Eq. 36–38 can also be derived from physical arguments which serve to reveal the origin of the expressions in Eq. 38. To this end, consider the rates of the various processes occurring in the repressor-free state and the pool of repressor-bound states. At steady state, transcription from the repressor-free state occurs at the rate

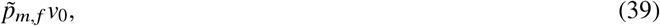

and transitions from the repressor-free to the repressor-bound state, which can occur by two mutually exclusive pathways, namely association of cytosolic repressor with *O*_1_ or *O*_2_, occur at the rate

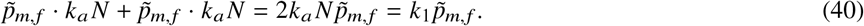

The corresponding expressions for the repressor-bound state follow from two properties of repressor-bound species that were obtained from perturbation theory, namely they are dominated by the looped species, i.e., *p_m,b_* ≈ *p*_*m*,12_, and satisfy detailed balance at steady state, i.e., 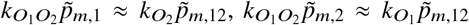. Hence, at steady state, transcription from the repressor-bound state occurs at the rate

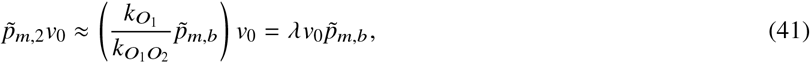

and transitions from the repressor-bound to the repressor-free state, which can occur by two mutually exclusive pathways, namely dissociation of the repressor from *O*_1_ · *R* or *O*_2_ · *R*, occur at the rate

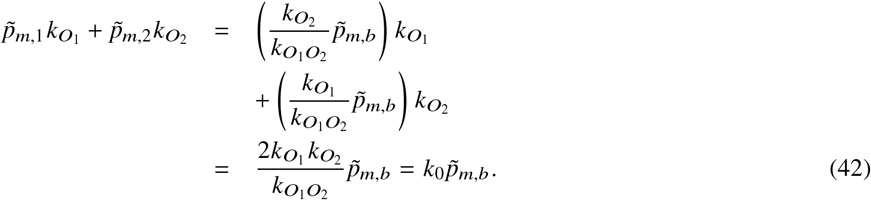

It follows from Eq. 39–42 that the master equations describing the steady state of the repressor-bound and repressor-free states are given by Eq. 36–38. More importantly, this approach yields physical insights into the origin of the expressions in Eq. 38. In Section 3.6, we shall exploit this physical approach to derive the leaky two-state model for the wild-type *lac* operon which has two auxiliary operators. Furthermore, in Appendix C, we show that more general sets of parameter values (e.g., where *k*_*O*1_ is of the same order as *k*_*O*_1_ *O*_2__) also result in the leaky two-state model.

### 3.4 The approximate solutions are close to the exact solution

It was argued above that the steady state of the full model can be approximated by the perturbation-theoretic solution to first-order, which in turn is approximated to order e by the steady state of the leaky two-state model. This assures us that the approximations are good for sufficiently small *ϵ*. However, it remains to verify that the particular value of *ϵ* of interest to us, determined by the value of *k*_*O*_1_ *O*_2__ in Table 1, is in fact sufficiently small for the approximations to be good.

We confirmed that the steady state marginal distribution of the full model is well approximated by the perturbation-theoretic solution to first order. It is shown in Appendix A that to first order, the generating function for the steady state marginal distribution of mRNA is

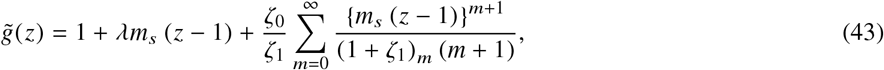

where (·)_*m*_ is the rising factorial, *m_s_* ≡ *ν*_0_/*d*_0_ is the mean number of mRNAs synthesized per mRNA lifetime in the unregulated operon (1, 2), and *ζ*_0_ ≡ *k*_0_/*d*_0_, *ζ*_1_ ≡ *k*_1_/*d*_0_ denote the mean number of transitions per unit mRNA lifetime between the repressor-bound and repressor-free states. We compared the steady state marginal mRNA distributions obtained from Eq. 43 with those obtained from simulations of the full model (Fig. 1) using the Optimized Direct Method implementation of Gillespie’s Stochastic Simulation Algorithm (10^6^ realizations let to run for 36000 s; initial state defined to be the looped operator state with no mRNAs present in the system) (34). Fig. 3a shows the steady state distributions calculated with the parameter values in Table 1 for the uninduced *lac* operon (obtained in the absence of the inducer). Under this condition, the approximate and simulated results agree well, but it remains to check if they also agree in the presence of the inducer. In principle, the inducer can bind to cytosolic repressor, which would decrease the association propensity *k_a_N*, as well as operator-bound repressor, which would increase the dissociation propensities *k*_*O*_1__, *k*_*O*_2__. However, since the inducer has a much higher affinity for the cytosolic repressor, only *k_a_N* will decrease in the presence of small inducer concentrations (13). Figs. 3b,c show that the steady state distributions obtained from simulations and perturbation theory agree well even when *k_a_N* is decreased 10- and 20-fold. The perturbation-theoretic solution is therefore valid for the uninduced and partially induced *lac* operon.

**Figure 3:**
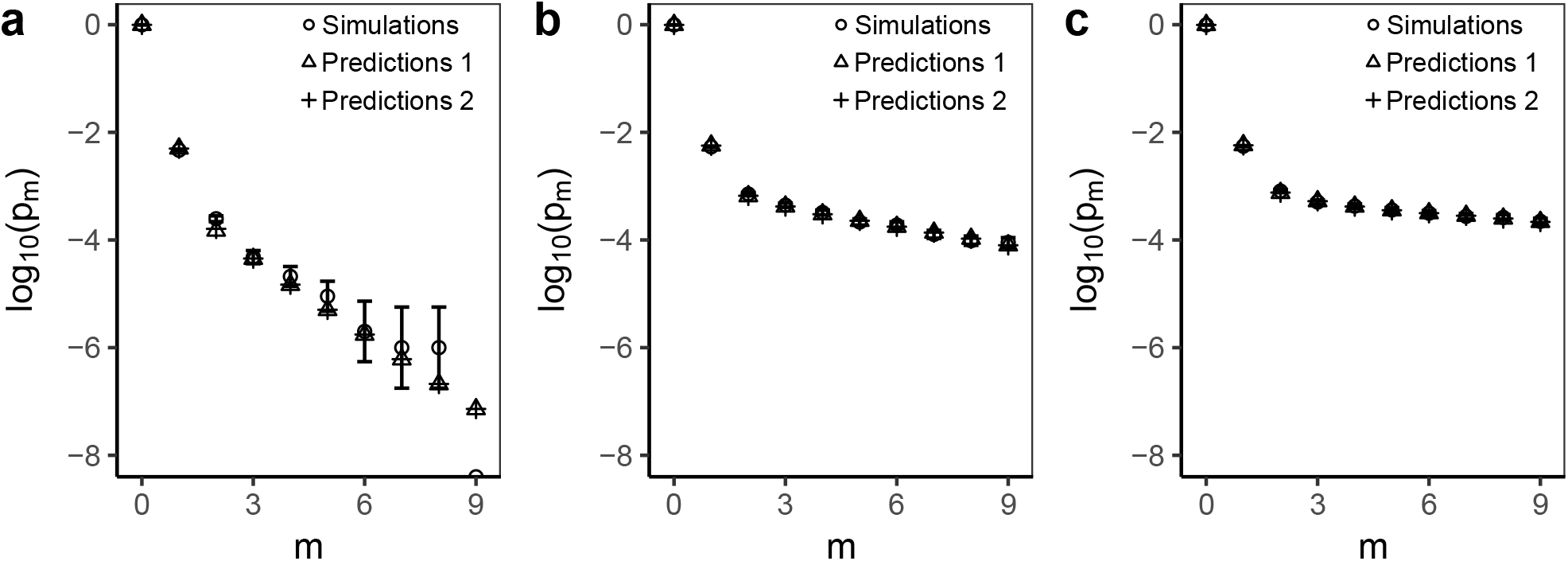
The steady state mRNA distributions obtained from simulations agree with predictions obtained from perturbation theory (labelled ‘Predictions 1’; see Eq. 43) and the leaky two-state model (labelled ‘Predictions 2’; see Eq. 44): The kinetic scheme in Fig. 1 was simulated for the parameter values (a) in the absence of the inducer (Table 1) and (b, c) the presence of inducer (Table 1 with *k_a_N* reduced 10-fold and 20-fold in (b) and (c), respectively). Error bars represent 95% binomial confidence intervals (obtained using Wilson method) around the observed frequency of a particular mRNA count in simulations.

The steady state marginal distribution obtained from the perturbation-theoretic solution to first order is essentially identical to the one obtained from the leaky two-state model (Fig. 3). Their equivalence is easily understood by comparing their generating functions. Indeed, it is shown in Appendix B that the generating function of the steady state marginal distribution for the leaky two-state model is

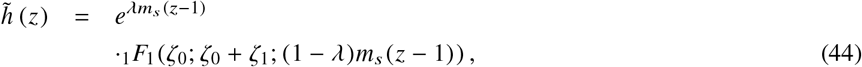

where _1_*F*_1_ (;;) denotes Kummer’s hypergeometric function of the first kind. This generating function is essentially equivalent to the generating function (Eq. 43) obtained from perturbation theory. Indeed, since *λ* ≪ 1, Eq. 44 can be rewritten as

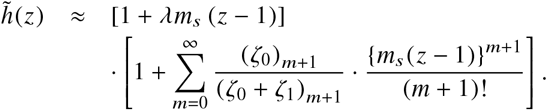

However, since *ζ*_0_ ≪ 1, *ζ*_1_, we have (*ζ*_0_)_*m*+1_ ≈ *ζ*_0_*m*! and (*ζ*_0_ + *ζ*_1_)_*m*+1_ ≈ *ζ*_1_ (*ζ*_1_ + 1)_*m*_, which imply that 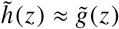, i.e, the generating functions of the steady state mRNA distributions obtained from the leaky two-state model and perturbation theory are essentially identical.

Taken together, the foregoing results imply that in the uninduced and partially induced *lac* operon, the steady state marginal mRNA distribution of the full model is well approximated by that obtained from the leaky two-state model.

### 3.5 The leaky two-state model yields simple expressions and physical insights

It is shown in Appendix B that the steady state marginal probability of the leaky two-state model 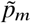 can be derived from the generating function (Eq. 44). However, we gain more physical insight by confining our attention to the latter. Indeed, the generating function (Eq. 44) is the convolution of Poisson and negative hypergeometric distributions, which shows that the steady state mRNA distribution is derived from two different sub-populations of mRNA. Indeed, since λ is the marginal probability of the state *O*_2_ · *R*, the Poissonian function represents the mRNA produced by transcription from the operon state *O*_2_ · *R*. On the other hand, since *λ* ≪ 1, the hypergeometric function is essentially identical to the generating function for the two-state model (5) and represents the mRNA produced by transcription from the free state *O*. We show below that although λ is small, transcription from *O*_2_ · *R* cannot be ignored because it provides most of the mRNA in the uninduced *lac* operon.

It follows from Eq. 44 that the mean *μ*(*m*) and variance *σ*^2^(*m*) of the marginal mRNA distribution are

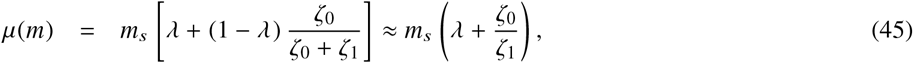

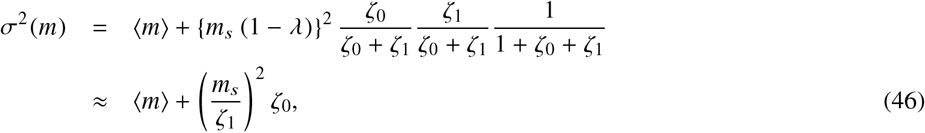

where the approximate results are obtained by appealing to the relations *λ* ≪ 1 and *ζ*_0_ ≪ 1 ≪ *ζ*_1_. Since *λ* and *ζ*_0_/(*ζ*_0_+*ζ*_1_) ≈ *ζ*_0_/*ζ*_1_ are the marginal probabilities of the states *O*_2_ · *R* and *O*, Eq. 45 shows that the mean consists of mRNA obtained by transcription from both *O*_2_ · *R* and O. However, in the uninduced *lac* operon, most of the mRNA is obtained by transcription from *O*_2_ · *R* since the parameter values in Table 1 imply that in the absence of inducer, *λ* = 4 × 10^−4^ is very small, but *ζ*_0_/*ζ*_1_ = 1 × 10^−4^ is even smaller. This result is consistent with the data of Choi et al. who observed that in uninduced cells, 80 % of the LacY molecules were derived from transcription of *O*_2_ · *R* (13). Eq. 46 shows that the variance also contains two terms, but only the second term reflects the variance due to regulation since the first term appears even in one-state models which have no regulation (1). Moreover, the variance due to regulation is entirely due to transcription from the repressor-free state *O*. We show below that transcription from *O*_2_ · *R* produces mRNA bursts that are so short-lived that they are averaged out on the slow time scale.

Although Eq. 45–46 provide simple expressions for the mean and variance, noisy gene expression is often characterized in terms of the size and frequency of transcriptional (mRNA) and translational (protein) bursts. In particular, Choi *et al.* suggested that in the *lac* operon, transcriptional bursts can arise from two distinct types of repressor dissociation (13):

1. A *partial* dissociation occurs when a repressor in the looped state dissociates from *O*_1_ but not *O*_2_, thus leading to the state *O*_2_ · *R* which allows transcription. However, since the repressor is still in the neighborhood of *O*_1_, it usually rebinds to *O*_1_ rapidly, and we expect only a *small* transcriptional burst. In other words, partial dissociations lead to transcription from the operon state *O*_2_ · *R*, and could result in small transcriptional bursts. These bursts are so short-lived that they are unlikely to yield more than one mRNA, but whenever this single mRNA is produced, it is likely to yield small *protein* bursts.
2. A *complete* dissociation occurs when the repressor dissociates from both operators and enters the cytosol. In this case, the repressor may take a relatively long time to rebind to *O*_1_, thus allowing a *large* transcriptional burst. Thus, complete dissociations lead to transcription from the operon state *O*, and yield large transcriptional bursts.

It is therefore convenient that Eq. 45–46 can be rewritten in terms of the size and frequency of the small and large transcriptional bursts. To see this, observe that during partial dissociations, *O*_2_ is repressor-free for the time period 1/k_*O*_1_*O*_2__, so that the mean size of the small transcriptional bursts is

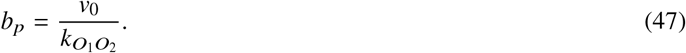

On the other hand, detailed balance at steady state implies that the mean frequency of partial dissociations per cell cycle is

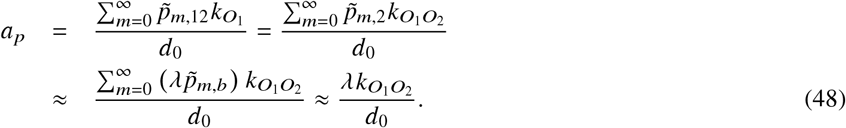

It follows that *a_p_b_p_* equals *m_s_ λ*, the first term of Eq. 45. Similarly, during complete dissociations, the operon is repressor-free for the time period 1/*k*_1_, so that the mean size of large transcriptional bursts is

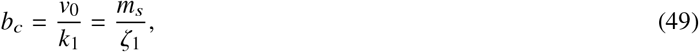

and the mean frequency of the complete dissociations per cell cycle is

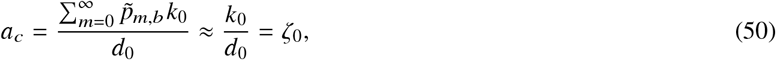

which implies that *a_c_b_c_* equals *m_s_ζ*_0_/*ζ*_1_, the second term of Eq. 45, and 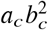 equals *ζ*_0_ (*m_s_/ζ*_1_)^2^, the second term of Eq. 46. It 2 follows that Eq. 45–46 can be rewritten as

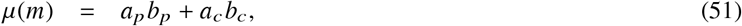

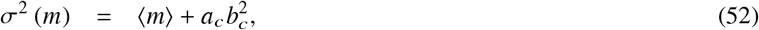

These equations show that partial dissociations contribute to the mean, but not the variance, because the resultant small transcriptional bursts, which occur on the fast time scale 1/*k*_*O*_1_O_2__, are averaged out on the slow time scale.

### 3.6 Steady states of the wild-type *lac* operon also follows the leaky two-state model

It was shown in Section 3.3 that the leaky two-state model corresponding to the simplified model with only one auxiliary operator (Fig. 1) could also be derived by appealing to two physical properties of the repressor-bound states at steady state, namely they satisfy detailed balance and are dominated by the looped states. Now the wild-type *lac* operon contains two auxiliary operators *O*_2_ and *O*_3_, which results in two additional significant states of the operon, namely *O*_3_ · *R* and the looped state *O*_3_ · *R* · *O*_1_ (32, 33). We shall now appeal to the foregoing physical properties of repressor-bound states to show that at steady state, the wild-type *lac* operon also follows Eq. 36–37, but *λ, k*_0_, and *k*_1_ are now given by the expressions

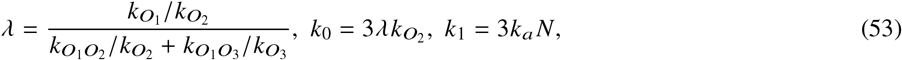

where *k*_*O*_3__ denotes the propensity of repressor dissociation from *O*_3_, and *k*_*O*_1_*O*_3__ denotes the propensity of *O*_3_ · *R* · *O*_1_ loop formation. To see this, observe that at steady state, transcription from the repressor-free state occurs at the rate 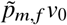, and transitions from the repressor-free to the repressor-bound state occur at the rate

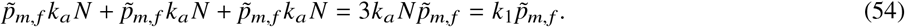

To determine the corresponding rates for the repressor-bound state, note that the repressor-bound states are predominantly in the looped state, i.e.,

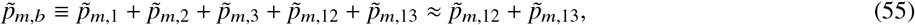

and satisfy the principle of detailed balance, i.e.,

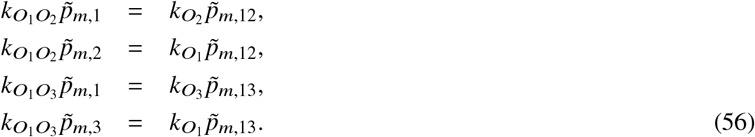

Eq. 55–56 can be solved to obtain

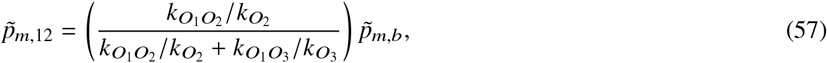

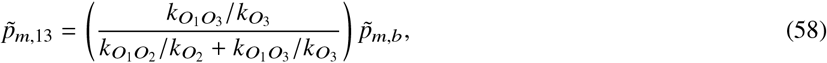

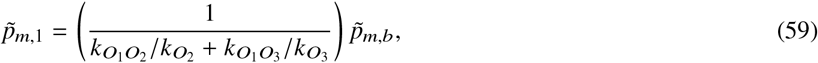

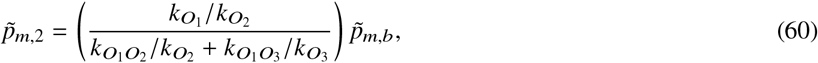

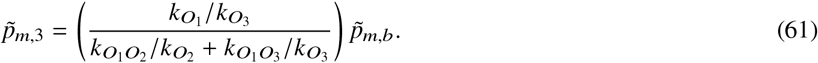

It follows from Eq. 59–61 that transcription from the repressor-bound state occurs at the rate

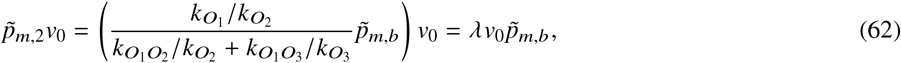

and transitions from the repressor-bound to the repressor-free state occur at the rate

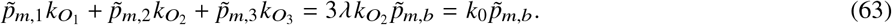

These rate expressions imply that at steady state, the master equations describing the interactions between the repressor-bound and repressor-free states of the wild-type *lac* operon are also given by Eq. 36–37, albeit with different parameter values given by Eq. 53. This result provides rigorous justification for our earlier work concerned with the steady state protein distributions in the wild-type *lac* operon (27). Moreover, it shows that the two physical facts inferred from perturbation theory, namely repressor-bound states are dominated by the looped state and satisfy detailed balance, are sufficient for extending the model to more complex scenarios.

## 4 CONCLUSIONS

Intuition suggests that due to fast DNA looping, the repressor-bound states of the *lac* operon are predominantly in the looped state and satisfy detailed balance. In earlier work, we derived these results by a heuristic approach, and showed that the resulting steady state protein distributions were in good agreement with stochastic simulations of the full model (27). We have shown above that singular perturbation theory provides a rigorous justification for the foregoing approximations. More precisely, application of perturbation theory shows that at steady state

1. To zeroth order, the operon is always in a looped state; in particular, the probabilities of the free and non-looped repressor-bound states are zero.
2. To first order, the probabilities of the repressor-bound states are in detailed balance, and the dynamics reduce to those of the leaky two-state model, a variant of the Peccoud-Ycart two state-model in which the bound state also permits transcription at a small, but significant, rate.

The leaky two-state model equations can be easily solved to obtain simple and physically meaningful expressions for the mean, variance, and the generating function of the mRNA distribution. Finally, the physics inferred from perturbation theory can be also used to derive the reduced (leaky two-state) model equations for more complex regulatory architectures which obviates the need for going through the intricate procedures of perturbation theory. This is a useful tool that is likely to facilitate the analysis of complex mechanistic models with auxiliary operators and DNA looping.

## A STEADY STATE SOLUTION OF FULL MODEL TO FIRST ORDER

Since we wish to derive the steady state marginal probability *p_m_* ≡ *p_m,b_* + *p_m,f_*, it is convenient to replace either *p_m,b_* or *p_m,f_* with *p_m_*. We have chosen to replace *p_m,b_* with *p_m_*, i.e., we shall solve for the zeroth- and first-order approximations of *p_m_* and *p_m,f_* (instead of *p_m,b_* and *p_m,f_*) at steady state. If *p_m_* has the perturbation expansion 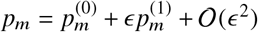, the perturbation expansions of *p_m,b_* and *p_m,f_* imply that 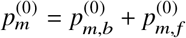 and 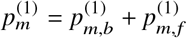.

### A.1 Zeroth-order solution

Adding Eq.19–20 yields

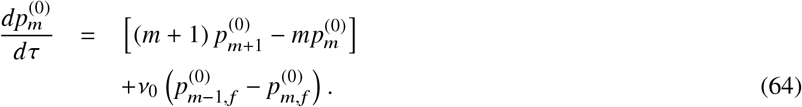

We shall obtain steady state solutions of Eq.20–21 and Eq.64 by the method of generating functions. To this end, let 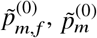 denote the steady states of these equations, and let 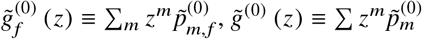 denote the corresponding generating functions. Multiplying both sides of Eq.20–21 and Eq.64 with *z^m^* and summing over *m* gives at steady state

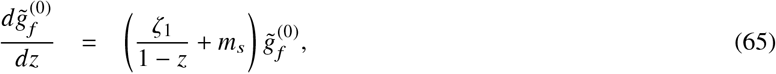

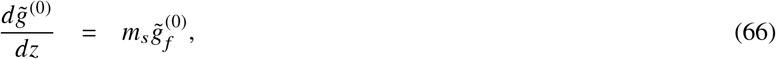

where 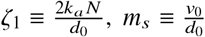 and, the boundary condition

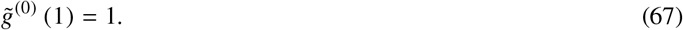

To solve these equations, note that integrating Eq.65 yields 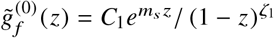 where *C*_1_ is a constant of integration. Since 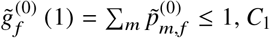 must be zero which implies that 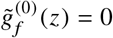. Then it follows from Eq.66–67 that

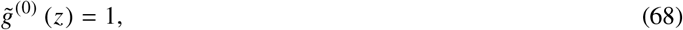

and 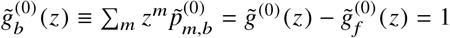. But 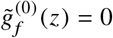 and 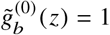 imply that to zeroth-order, the steady state probabilities are

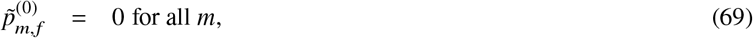

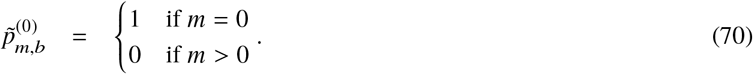

### A.2 First-order solution

To solve for the generating functions of 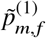 and 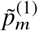, add Eq.30–31 to get

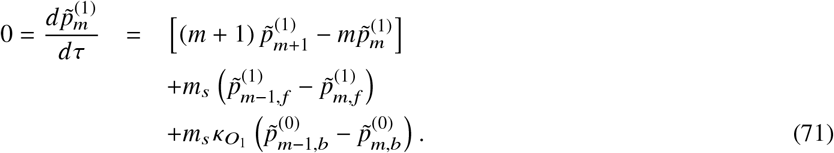

Now define 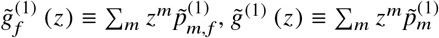, multiply both sides of Eq.31 and Eq.71 with *z^m^*, and summing over all values of *m* gives the equations

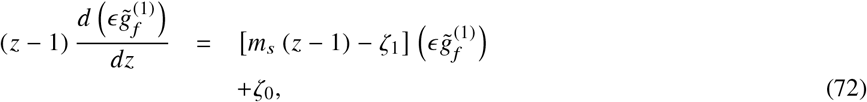

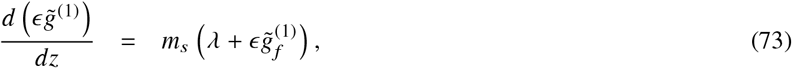

where 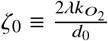 and, the boundary condition

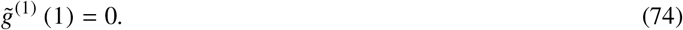

Solving Eq.72 under the constraint of bounded 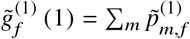 yields

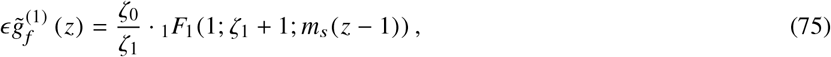

which implies that

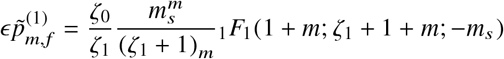

for all *m* ≥ 0. Now it follows from Eq.73–74 that

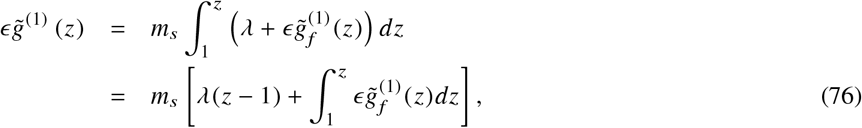

Substituting the series for the hypergeometric function in Eq.75, and integrating term-by-term yields

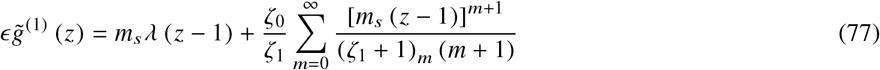

where (·)_*m*_ denotes the Pochhammer symbol (rising factorial). It follows from Eq.77 that

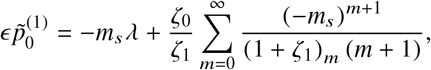

and Eq.76 implies that

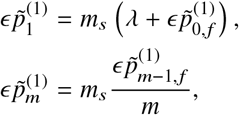

for all *m* ≥ 2. Given these expressions for 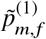 and 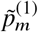, we can determine 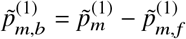.

It follows from Eq.68 and Eq.77 that to first order, the generating function for the steady state marginal distribution of mRNA is

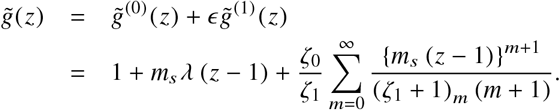

## B STEADY STATE SOLUTION OF LEAKY TWO-STATE MODEL

Once again, it is convenient to replace *p_m,b_* by the marginal probability *p_m_*≡*p_m,b_* + *p_m,f_*, which evolves in accordance with the equation

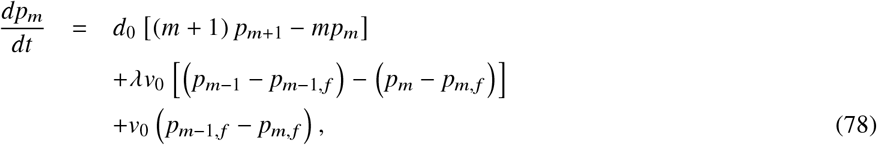

obtained by adding Eq.36–37. We shall solve for the steady states 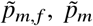 of Eq.37 and Eq.78. To this end, let 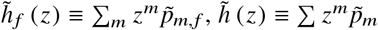, multiply both equations by *z^m^*, and sum over *m* to obtain at steady state

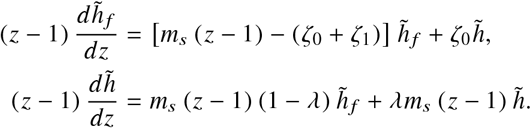

Moreover, since Σ_*m*_ *p_m_*= 1, 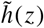 satisfies the boundary condition 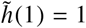. The above system of two first-order differential equations can be reduced to the single second-order differential equation

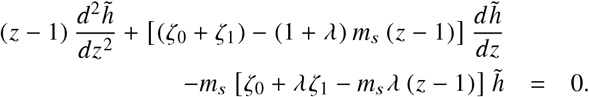

If we define *ν* =*m_s_* (1 − *λ*) (*z* − 1) and 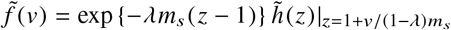, the above equation reduces to Kummer’s standard differential equation

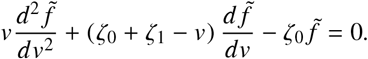

Since 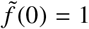, we obtain the solution 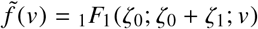, which can be rewritten as

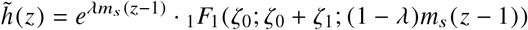

by returning to the the original variables. Differentiating this equation m times yields

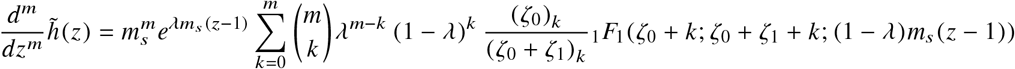

which implies that the steady state marginal distribution of mRNA is

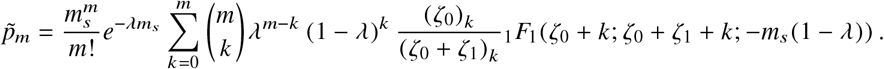

## C OTHER PARAMETER COMBINATIONS ALSO RESULT IN LEAKY TWO-STATE MODEL

Let us consider a hypothetical scenario where *k*_*O*_1__ is comparable to *k*_*O*_1_ *O*_2__. In this case, intuition suggests that the repressor-bound states are predominantly in the looped and *O*_2_· *R* states. Once again, we follow the perturbation-theoretic approach outlined above to derive reduced equations. Multiplying Eq.4 and Eq.8–10 with the redefined perturbation parameter 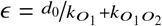, and defining 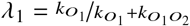 and 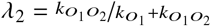 gives the slow equations

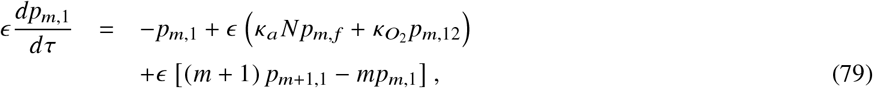

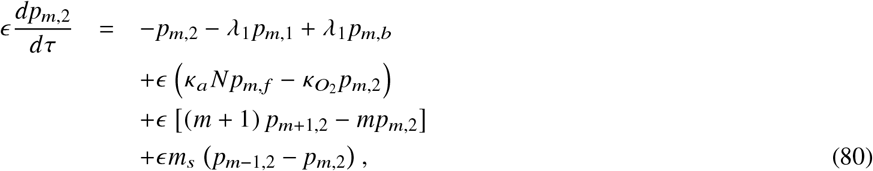

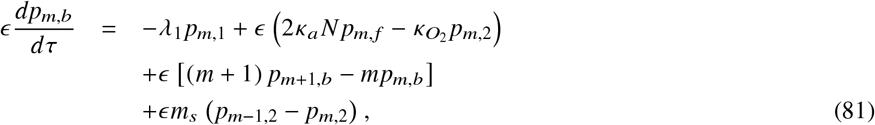

Next we substitute the power series expansions

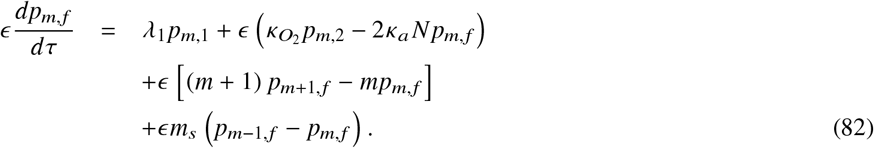

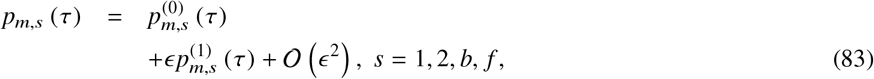

in Eq.79–82, and collect terms with coefficients *ϵ*^0^ and *ϵ*^1^ to determine the zeroth- and first-order relations as follows. The zeroth-order equations are

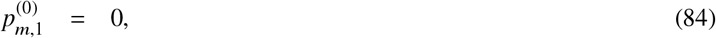

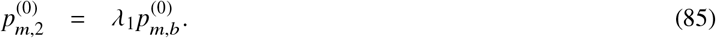

Note that in this case, the zeroth-order equations reveal a partially induced operon as expected. The first-order equations are

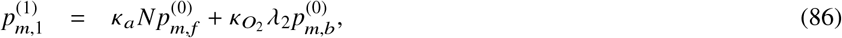

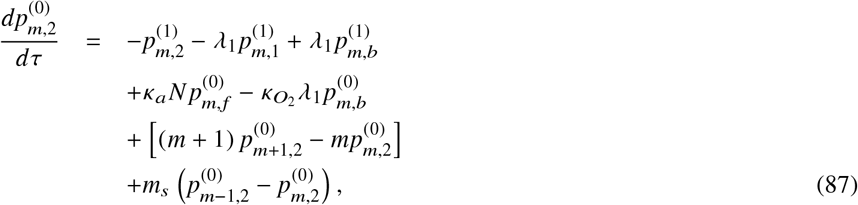

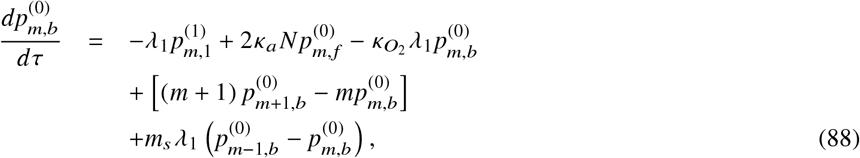

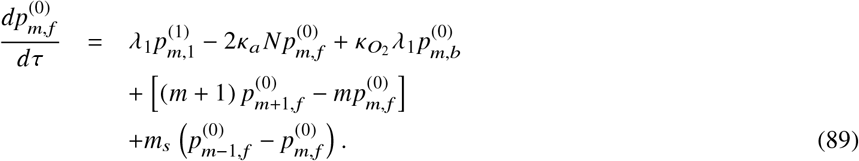

Using Eq.86 in Eq.88–89, at steady state

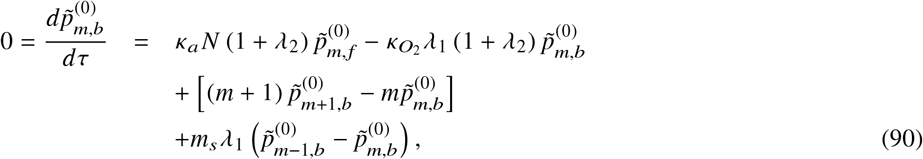

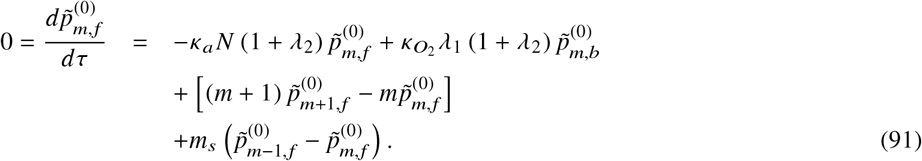

Above equations are formally identical to equations for a leaky two-state model and can be solved using the generating function method. Notably, in this case, the leaky-two state model is reflected in the zeroth-order terms due to partial induction of the operon.

